# Syntenin orchestrates matrix degradation by controlling MT1-MMP secretion in small extracellular vesicles and invadopodia formation

**DOI:** 10.1101/2025.09.01.673493

**Authors:** Marie Huber, Rania Ghossoub, Raphaël Leblanc, Guido David, Pascale Zimmermann, Sylvie Thuault

**Author notes:** Corresponding author: Sylvie Thuault.

## Abstract

Extracellular matrix (ECM) remodelling is essential for tumour progression. The metalloproteinase MT1-MMP promotes cell invasion by degrading ECM components. MT1-MMP accumulates at invadopodia, actin-rich structures mediating ECM degradation, and is also present on the surface of small extracellular vesicles (sEV), key players in cell-to-cell communication. The role of sEV-associated MT1-MMP in ECM degradation and the mechanisms supporting MT1-MMP loading into sEV are unknown. We previously established that the syntenin PDZ protein, in association with the syndecan (SDC), is essential for sEV from endosomal origin biogenesis. In the present study, we demonstrate that syntenin contributes to ECM degradation *via* invadopodia in MDA-MB-231 triple negative breast cancer (TNBC) cells, whereas SDC limit it. Moreover, we identify that syntenin colocalizes with MT1-MMP in late endosomes and directly interacts with MT1-MMP. Furthermore, we decipher that both syntenin and SDC are essential for the specific loading of MT1-MMP into sEV. Additionally, we show that sEV-mediated ECM degradation depends on MT1-MMP activity and syntenin. Our findings show how syntenin-SDC-MT1-MMP complexes orchestrate ECM degradation and pave the way for innovative rational approaches to control TNBC invasion.

## Introduction

Extracellular matrix (ECM) remodeling is critical in physiological and pathological conditions such as cancer. One key process underlying cancer cell invasiveness is the formation of invadopodia, specialized actin-rich protrusions of the plasma membrane that coordinate local ECM degradation (Eddy et al., 2017; Weaver, 2006). Invadopodia serve as platforms for the targeted delivery and activation of matrix-degrading enzymes, allowing cancer cells to penetrate surrounding stromal tissues and disseminate to distant sites. Among the proteolytic enzymes recruited to invadopodia, membrane-type 1 matrix metalloproteinase (MT1-MMP, or MMP14) emerges as the major player protease (Linder, 2007; Poincloux et al., 2009; Frittoli et al., 2011; Castro-Castro et al., 2016; Gifford and Itoh, 2019; Hey et al., 2022). MT1-MMP is able to degrade structural components of the ECM such as collagen, laminin, and fibronectin (Itoh, 2015), cleave adhesion proteins such as CD44 (Kajita et al., 2001), α_v_ integrins (Deryugina et al., 2002; Ratnikov et al., 2002) and syndecan (Endo et al., 2003), as well as activate other matrix metalloproteinases (MMP), such as pro-MMP-2 (Sato et al., 1994) and pro-MMP-13 (Knäuper et al., 1996). Consistent with its proteolytic capacity, MT1-MMP expression is frequently upregulated in aggressive cancers and strongly correlates with increased invasion, metastasis, and poor prognosis (Liu et al., 2025).

While the localization of MT1-MMP to invadopodia has been extensively studied, some studies have demonstrated that MT1-MMP is also present on the surface of extracellular vesicles (EV) released by cancer cells (Hakulinen et al., 2008; Clancy et al., 2015; Shimoda and Khokha, 2017). Among EV, exosomes, of endosomal origin, and microvesicles, derived from the plasma membrane, were historically distinguished (Lötvall et al., 2014). However, recent advances in the field (MISEV2023) recommended avoiding classification of EV based on their biogenesis, and instead propose size-based terminology, distinguishing small EV (sEV < 200nm, enriched in exosomes), from large EV (lEV > 200nm, enriched in microvesicles) (Welsh et al., 2024). EV are essential for intercellular communication in physiological and pathological conditions (van Niel et al., 2018). EV-associated MMP have been implicated in key processes of tumour progression, such as migration, angiogenesis, ECM remodelling and premetastatic niche formation (Shimoda and Khokha, 2017; Thuault et al., 2022). Nevertheless, the mechanisms regulating the incorporation of MT1-MMP into EV remain largely unknown, and the biological significance of this vesicle-associated pool of MT1-MMP has not been fully elucidated.

Importantly, the intracellular domain (ICD) of MT1-MMP contains a class III PDZ-binding motif (PDZ BM) (Thuault et al., 2022), which is conserved across species and mediates interactions with PDZ domain-containing proteins such as SNX27 (Sharma et al., 2019). Our team has demonstrated that syntenin, a multifunctional PDZ domain-containing adaptor protein, plays a pivotal role in endosomal trafficking (Latysheva et al., 2006; Zimmermann et al., 2005), and in the biogenesis of sEV from endosomal origin (also referred to as exosomes) (Baietti et al., 2012; Ghossoub et al., 2014). sEV from endosomal origin originate from early endosomes that mature into late endosomes, where inward budding of the limiting membrane generates multivesicular bodies (MVB) containing intraluminal vesicles (ILV). Upon fusion of MVB with the plasma membrane, ILV are released into the extracellular space as sEV from endosomal origin. The interactions of syntenin with the accessory protein ALG-2 interacting protein X (ALIX) of the endosomal sorting complex required for transport (ESCRT) machinery, and with syndecan (SDC) heparan sulfate proteoglycans, are essential for syntenin control of the biogenesis of sEV from endosomal origin (Baietti et al., 2012). In addition, syntenin is responsible for the recycling of SDC to the plasma membrane (Zimmermann et al., 2005). Both processes require a direct interaction between the PDZ BM of SDC and the two PDZ domains of syntenin (Grootjans et al., 2000). By this dual function, syntenin not only controls the biogenesis of sEV from endosomal origin, but also their molecular composition and uptake by recipient cells (Leblanc et al., 2020; Kashyap et al., 2021). Thus, syntenin influences the downstream biological activity of sEV. Elevated syntenin expression has been linked to poor clinical outcome in several cancer types (Das et al., 2012; Qian et al., 2013; Yang et al., 2013). In fact, syntenin has been implicated in key pro-tumoral processes including adhesion (Zimmermann et al., 2001), migration (Kashyap et al., 2015; Imjeti et al., 2017; Irmer et al., 2025), and epithelial-to-mesenchymal transition (EMT), notably through the regulation of the TGF-β/Smad signalling pathway (Hwangbo et al., 2016). However, its direct involvement in ECM degradation has not been established.

Given the role of syntenin in orchestrating the endosomal trafficking and the sorting of cargoes into sEV through its PDZ domains (Baietti et al., 2012; Latysheva et al., 2006; Leblanc et al., 2020), syntenin might be a critical player in the recruitment of MT1-MMP into the vesicular pathway. Thus, we hypothesize that the syntenin/SDC pathway may actively regulate the invasive potential of tumour cells by mediating the recycling of MT1-MMP for its incorporation into invadopodia and/or the selective sorting of MT1-MMP into sEV. Through these mechanisms, cancer cells may enhance their ECM-degradative capacity not only at the plasma membrane *via* invadopodia, but also in the extracellular space through sEV-mediated ECM degradation. This dual localization of MT1-MMP could provide a spatially extended mode of ECM remodelling, supporting both local invasion and the preparation of metastatic niches. Elucidating the molecular determinants governing MT1-MMP sorting into sEV, and the role of syntenin in this process, may therefore uncover critical mechanisms of tumour dissemination and reveal new therapeutic opportunities aimed at impairing cancer cell invasion.

In this study, we show that syntenin positively regulates ECM degradation *via* invadopodia in MDA-MB-231, TNBC-derived cells, while SDC limit this process. We show that syntenin colocalizes with MT1-MMP in late endosomes, the cellular compartment from which sEV from endosomal origin emerge. In addition, we identify that syntenin interacts directly with the ICD of MT1-MMP *via* its PRR motif. Furthermore, we demonstrate that syntenin and SDC are essential determinants of the specific loading of MT1-MMP into sEV without affecting either the cell surface levels or the endocytosis of MT1-MMP. Finally, we demonstrate that sEV participate in ECM degradation through MT1-MMP activity, a syntenin-dependent process.

## Results

### Syntenin positively regulates extracellular matrix degradation *via* invadopodia

We investigated the contribution of syntenin and SDC in the ability of the invasive MDA-MB-231 TNBC-derived cells to form invadopodia and degrade the ECM. To this end, MDA-MB-231 cells were transfected either with a non-targeting siRNA (siCTRL) or siRNAs targeting syntenin (siSYNT_03 and siSYNT_04) and SDC 1 and 4 (siSDC1+4), the two SDC the most expressed in these cells (Beauvais and Rapraeger, 2003). The cells were then seeded on a fluorescently-labelled gelatin artificial ECM for 5 hours (Figure 1A) allowing to detect ECM degradation, which appeared as dark, non-fluorescent areas. Invadopodia were identified by co-localization of cortactin and TKS5, two core invadopodia components (Eddy et al., 2017), while mature, enzymatically active invadopodia were identified by the co-localization of TKS5, cortactin and ECM degradation.

**Figure 1:**
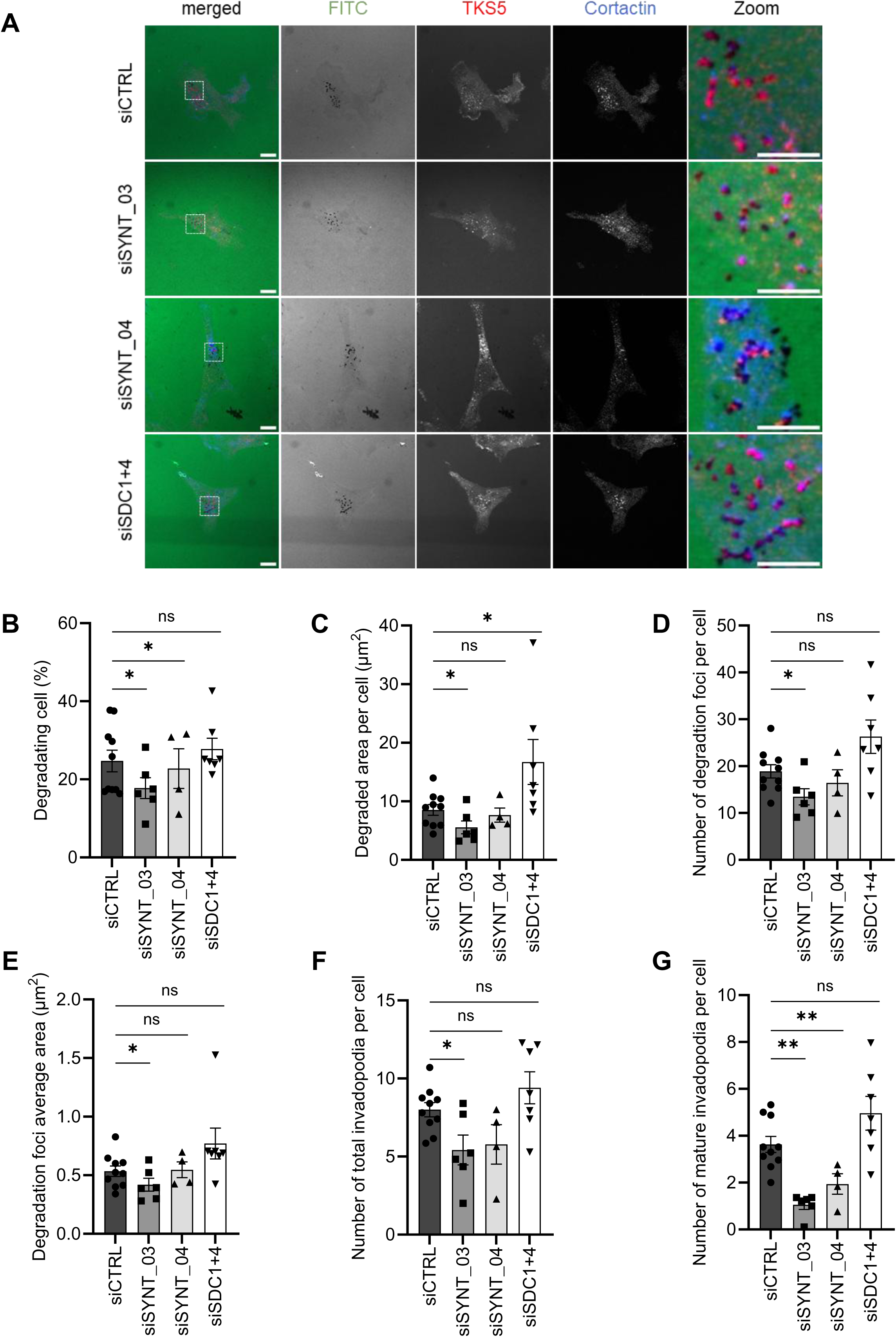
Syntenin positively regulates ECM degradation *via* invadopodia in MDA-MB-231 TNBC cells, while SDC limit it. (**A**) Representative images of MDA-MB-231 cells transfected with a non-targeting siRNA (siCTRL), siRNA targeting syntenin (siSYNT_03 and siSYNT_04), or siRNA targeting SDC1 and SDC4 (siSDC1+4), seeded on fluorescently-labeled gelatin (FITC-gelatin) for 5 hours. Cells were fixed and stained with anti-TKS5 and -cortactin antibodies to identify invadopodia. Dark spots in the FITC-gelatin correspond to matrix degradation. The white-boxed regions are enlarged (zoom). Scale bars represent 10 μm in non-enlarged images, 5 µm in enlarged images. (**B-G**) The ability of MDA-MB-231 cells to degrade fluorescently labeled gelatin was quantified: (**B**) the percentage of degrading cells, (**C**) the degraded area per cell, (**D**) the number of degradation foci per cell, (**E**) the averaged area of each degradation foci, the number (**F**) of invadopodia and (**G**) of mature invadopodia per cell. Dot plots represent the mean values ± SEM of ten independent experiments for siCTRL, six for siSYNT_03, four for siSYNT_04 and seven siSDC1+4. The different parameters of matrix degradation were quantified using a custom Fiji macro (for the percentage of degrading cell: 10 random fields per condition per experiment, for the other parameters: 25 cells per condition per experiment). The Paired Student’s t-test with Welsh correction was used for statistical analysis. *** p ≤ 0.001, ** p ≤ 0.01, * p ≤ 0.05, ns: not significant.

Silencing syntenin using two independent siRNA sequences (siSYNT_03 and siSYNT_04) led to a significant reduction in ECM degradation, as evidenced by a decrease in the percentage of cells exhibiting ECM degradation activity (Figure 1B). Further quantitative analysis revealed also a significant decrease in the total area of degraded ECM per cell following syntenin silencing *via* siSYNT_03 (Figure 1C). This reduction can be attributed to both a lower number of degradation foci per cell (Figure 1D), and a decrease in the mean size of individual degradation foci (Figure 1E) indicating an overall impairment in ECM remodelling. Similar trends were observed with siSYNT_04 for these parameters, although they did not reach statistical significance. A reduction in the number of invadopodia was observed, reaching significance with siSYNT_03 and showing a similar trend with siSYNT_04 (Figure 1F). Interestingly, this was accompanied by a more pronounced decrease in mature invadopodia (Figure 1G), suggesting that the reduction in ECM degradation induced by syntenin inhibition primarily reflects an impairment in MMP recruitment at precursor invadopodia.

In contrast, SDC depletion was associated with an increase in the total degraded area per cell (Figure 1C), without affecting the percentage of cells exhibiting ECM degradation activity (Figure 1B). However, SDC depletion did not affect the number of degradation foci (Figure 1D) or their size (Figure 1E), nor the number of total invadopodia (Figure 1F) or mature invadopodia (Figure 1G), although slight upward trends were observed for these parameters. These observations suggest that SDC do not regulate invadopodia formation/maturation *per se* but rather control their dynamics of assembly and di-assembly, or spatial organization. This deserves further investigation.

Together, these results reveal that syntenin influences positively ECM degradation by cells through its impact on invadopodia formation and activity, while SDC seem to limit the extent of ECM degradation.

### Syntenin colocalizes with MT1-MMP preferentially in late endosomes

Considering the impact of syntenin depletion on mature invadopodia, we next investigated whether syntenin colocalizes with the key invadopodia-associated protease MT1-MMP. Confocal microscopy imaging revealed a striking colocalization of endogenous syntenin and endogenous MT1-MMP within intracellular vesicular structures in MDA-MB-231 cells (Figure 2A). A Pearson’s correlation coefficient around 0.7 confirmed a significant overlap between syntenin and MT1-MMP signals (Figure 2B).

**Figure 2:**
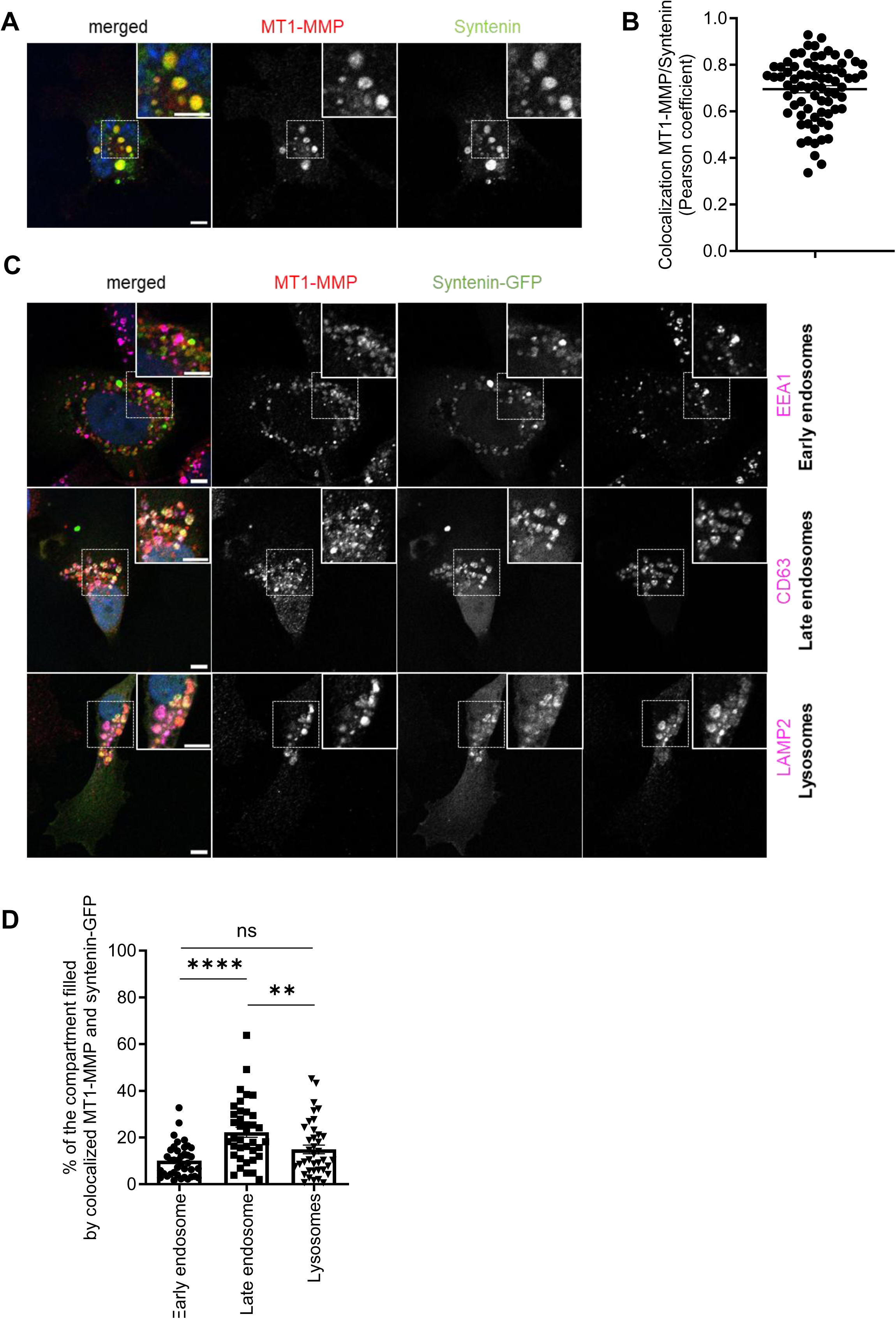
Syntenin colocalizes with MT1-MMP preferentially in late endosomes in MDA-MB-231 cells. (**A**) Representative confocal images showing the steady-state distributions of endogenous syntenin and endogenous MT1-MMP in MDA-MB-231 cells. In the merged image, the nucleus is stained with DAPI (blue), syntenin appears in green, and MT1-MMP in red. Scale bars represent 5 μm. (**B**) Pearson correlation coefficient for syntenin and MT1-MMP colocalization, calculated using the JACoP plugin in ImageJ. Dot plot represents Pearson coefficient of 75 cells ± SEM from three independent experiments (25 cells per experiment). (**C**) Representative confocal images of MDA-MB-231 cells expressing syntenin-GFP (green) stained for endogenous MT1-MMP (red) and for specific intracellular compartment (magenta): EEA1 for early endosomes, CD63 for late endosomes and LAMP2 for lysosomes. In the merged images, the nuclei are stained with DAPI (blue). Scale bars represent 5 μm. (**D**) Dot plot represents the average percentage of each compartment filled by colocalized MT1-MMP and syntenin-GFP quantified in 40 cells ± SEM from two independent experiments (20 cells per condition per experiment), analysed using ImageJ. The Mann-Whitney test was used for statistical analysis. **** p ≤ 0.0001, *** p ≤ 0.001, ** p ≤ 0.01, * p ≤ 0.05, ns: not significant.

MT1-MMP has been reported to localize at various cellular sites, including the plasma membrane as well as within intracellular compartments, such as late endosomes (Poincloux et al., 2009, Planchon et al., 2018; Palmulli et al., 2025). Syntenin is found in multiple cellular compartments, most notably in the cytoplasm and the endosomal system (Baietti et al., 2012). To identify the specific cellular compartments where syntenin and MT1-MMP colocalize, MDA-MB-231 cells were transfected with syntenin-GFP, while endogenous MT1-MMP and various endogenous cellular compartments were labelled using specific antibodies (Figure 2C). Our results demonstrate that syntenin-GFP and endogenous MT1-MMP colocalize within multiple cellular compartments (Figure 2C). After analyzing images based on object segmentation, the most prominent colocalization was observed within late endosomes, as evidenced by their strong colocalization in the CD63-positive compartment. However, colocalization was also noted in early endosomes (EEA1-positive) and lysosomes (LAMP2-positive) (Figure 2D).

These observations suggest a functional association between syntenin and MT1-MMP and highlight the potential role of syntenin as a regulatory factor of MT1-MMP trafficking. In particular, syntenin could influence the sorting of MT1-MMP within the late endosomal compartment, a key site for vesicle sorting decisions.

### Syntenin directly interacts with MT1-MMP *via* its PRR motif

As previously mentioned, the ICD of MT1-MMP contains a class III PDZ BM, which, in light of its co-localization with syntenin and the role of syntenin in sEV cargo sorting *via* its PDZ domains, likely contributes to MT1-MMP recruitment into the vesicular pathway. Based on this, we investigated the interaction between MT1-MMP and syntenin by surface plasmon resonance (SPR) using the BIAcore technology. Biotinylated peptides corresponding to the wild-type intracellular domain of MT1-MMP (MT1-MMP ICD WT), to the last 19-amino-acids of the wild-type intracellular domain of SDC2 (SDC2 ICD WT), used as a positive control, and a mutant version lacking its PDZ BM essential for syntenin binding (Grootjans et al., 1997) (SDC2 ICD ΔFYA), used as a negative control, were immobilized on a streptavidin-coated sensor chip (Figure 3A). These immobilized peptides were then exposed to the PDZ domains of syntenin and its C-terminal extremity fused to GST (GST-Syntenin PDZ Tandem WT + Cter) at 10 µM (Figure 3B) for 300 s (association phase), followed by a 120 s of dissociation. Our results demonstrate that syntenin and MT1-MMP interact directly, although this interaction is weaker than that observed between syntenin and SDC2, one of its major binding partners (Grootjans et al., 1997). No interaction was observed with the negative control (SDC2 ΔFYA), confirming the specificity of the binding (Figure 3C). Kinetic analysis using increasing concentrations of purified GST-Syntenin PDZ Tandem WT + Cter (0 µM – 80 µM) confirmed the specificity of the syntenin/MT1-MMP interaction, as the binding response increased proportionally to the analyte concentration (Figure 3D).

**Figure 3:**
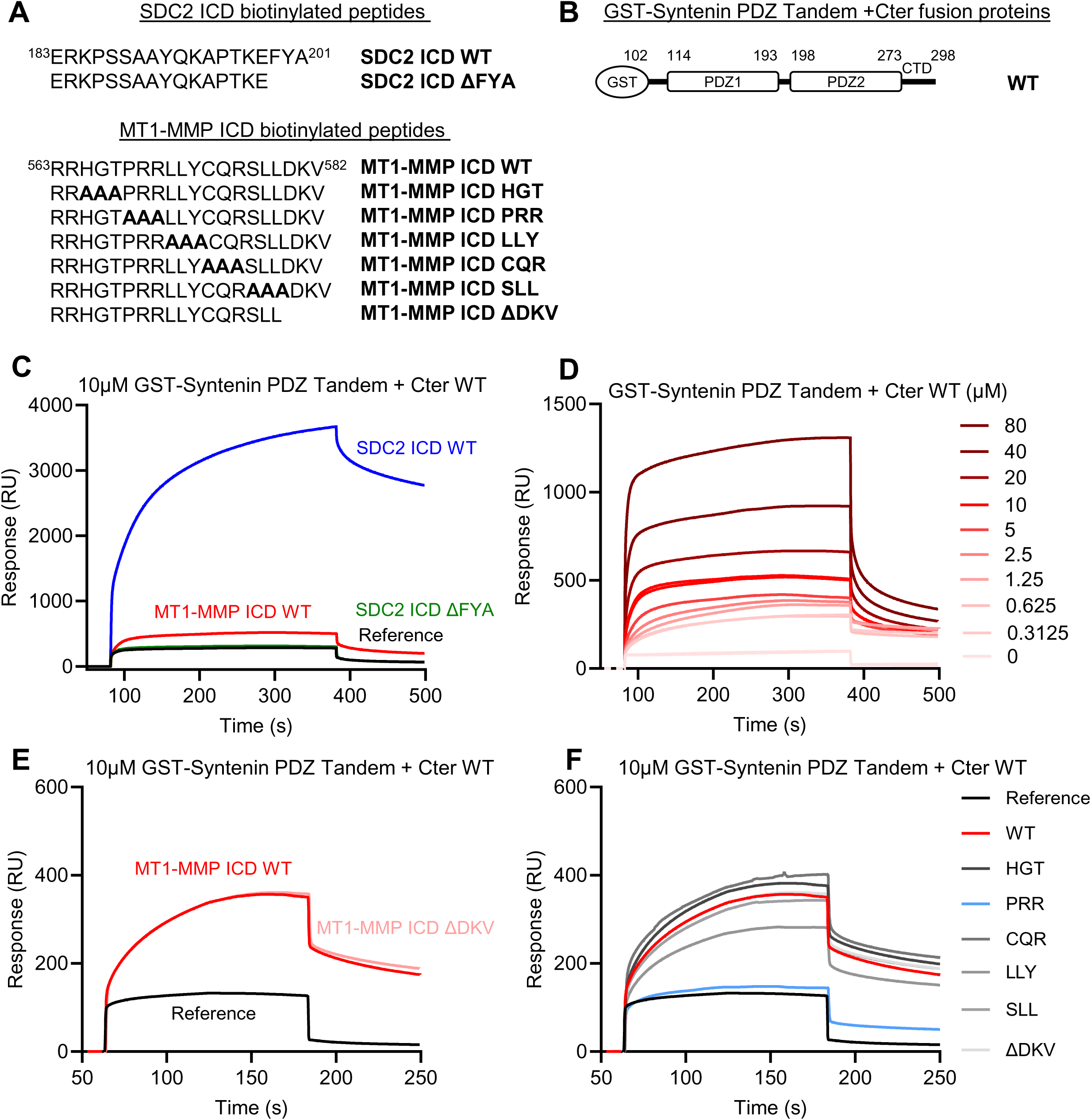
Syntenin directly interacts with MT1-MMP *via* the PRR motif. (**A**) Schematic representation of biotinylated peptides corresponding to the intracellular domain (ICD) of SDC2 and MT1-MMP, including their mutated and PDZ binding motif-depleted variants (mutated residues are in bold). (**B**) Schematic representation of GST-syntenin PDZ tandem fusion protein. (**C**) Biotinylated peptides corresponding to the ICD of MT1-MMP, SDC2 (positive control), and SDC2 lacking its PDZ binding motif (ΔFYA, negative control) were immobilized (1,000 RU) on a streptavidin-coated sensor chip followed by injection of GST-syntenin PDZ tandem WT (10 µM) for 300 s, followed by 120 s of dissociation. Representative sensorgram from at least three independent experiments is shown. (**D**) Biotinylated peptides corresponding to the ICD of MT1-MMP were immobilized (1,000 RU), and increasing concentrations of GST-syntenin PDZ tandem WT were injected for 300 s, followed by 120 s of dissociation. Representative sensorgram from at least three independent experiments is presented. (**E**) Biotinylated peptides corresponding to the ICD of MT1-MMP and its PDZ binding motif-depleted variant were immobilized (1,000 RU), followed by injection of GST-syntenin PDZ tandem WT (10 µM) for 120 s, with 120 s of dissociation. Representative sensorgram of N=1 chip, N=2 GST-syntenin productions is shown. (**F**) Biotinylated peptides corresponding to the ICD of MT1-MMP and its mutated variants (as described in (**A**)) were immobilized (1,000 RU), followed by injection of GST-syntenin PDZ tandem WT (10 µM) for 120 s, followed by 120 s of dissociation. Representative sensorgram of N=1 chip, N=2 GST-syntenin productions is presented.

We next examined whether their interaction is mediated by the PDZ BM of MT1-MMP. To test this hypothesis, a biotinylated peptide corresponding to the ICD of MT1-MMP lacking its PDZ BM (MT1-MMP ICD ΔDKV) (Figure 3A) was immobilized on a streptavidin-coated sensor chip, followed by injection of GST-Syntenin PDZ Tandem WT + Cter (10 µM) (Figure 3B) for 120 s association phase, followed by 120 s dissociation phase. The results indicate that the absence of the PDZ BM does not affect the interaction of MT1-MMP with syntenin PDZ Tandem domains (Figure 3E).

To identify the interaction motif within MT1-MMP, an alanine scanning mutagenesis approach was performed on the ICD of MT1-MMP (Figure 3A). Interestingly, alanine substitutions of the PRR motif of MT1-MMP completely abolished the interaction with syntenin, whereas alanine mutations in the other motifs did not alter the interaction (Figure 3F), indicating that the PRR motif of MT1-MMP is essential for syntenin recognition.

In conclusion, our SPR analysis demonstrates that syntenin directly interacts with the ICD of MT1-MMP and that this interaction does not involve the PDZ BM of MT1-MMP. We identified the PRR motif of MT1-MMP as essential for syntenin recognition. These findings provide new insights into the molecular basis of syntenin/MT1-MMP interaction and its potential functional implication, especially considering their co-localization in late endosomes.

### Syntenin and SDC are essential determinants of the specific loading of MT1-MMP into sEV

To determine whether syntenin and SDC influence MT1-MMP cellular trafficking, we examined their impact on MT1-MMP cell surface levels and endocytic uptake. For this, MDA-MB-231 TNBC cells were transfected with either a non-targeting siRNA (siCTRL) or siRNAs targeting syntenin (siSYNT_03 and siSYNT_04) and SDC 1 and 4 (siSDC1+4). Biotinylation of cell surface proteins followed by their analysis by Western blot revealed no significant changes in MT1-MMP abundance at the plasma membrane upon silencing of syntenin or SDC (Figure 4A-B). Moreover, tracking MT1-MMP internalization over 30 minutes showed comparable endocytosis rates between control and syntenin- or SDC-depleted cells (Figure 4C). These findings suggest that neither syntenin nor SDC affect MT1-MMP trafficking to the plasma membrane or its internalization.

**Figure 4:**
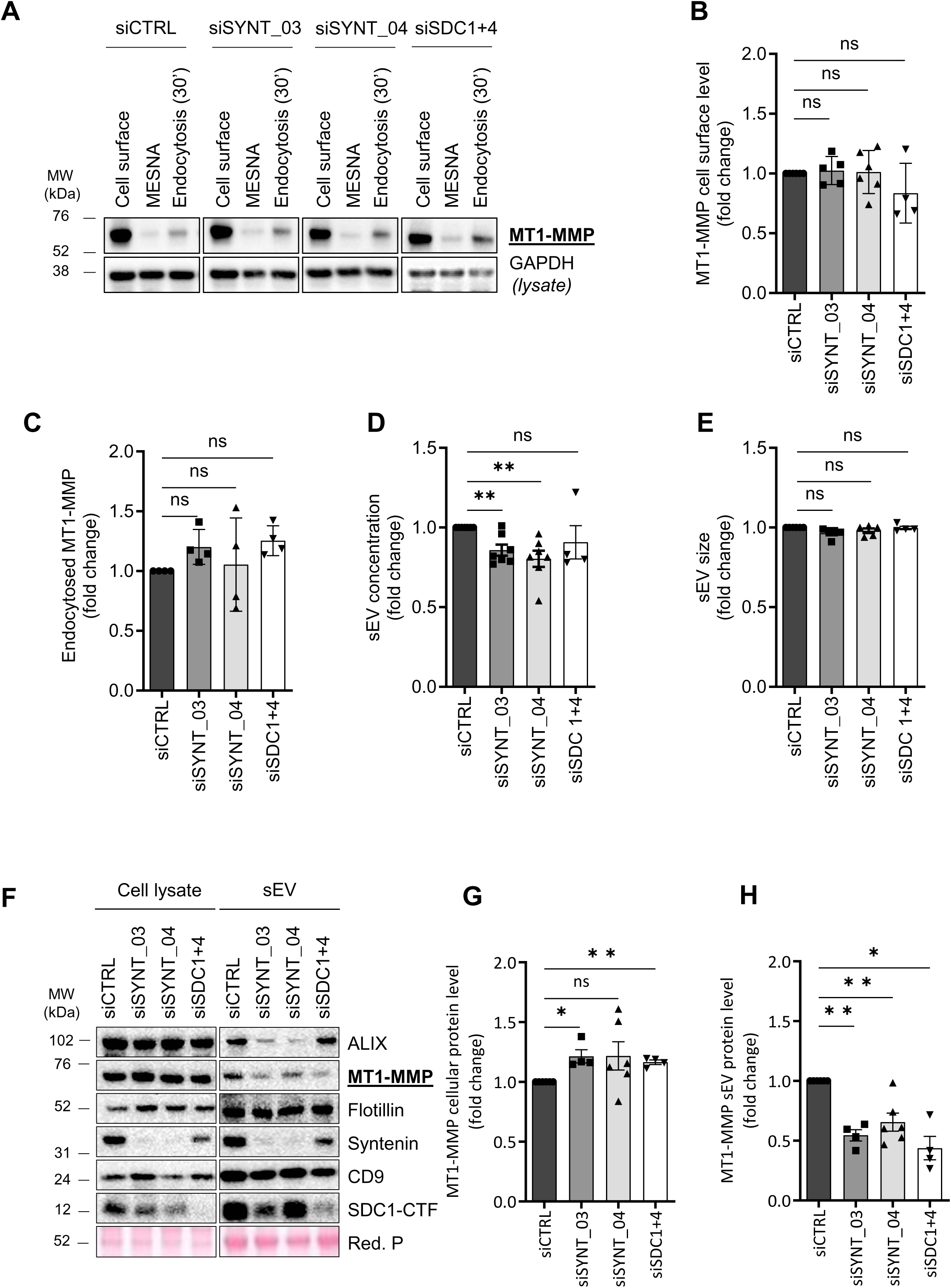
Syntenin and syndecans are essential determinants of the specific loading of MT1-MMP into sEV. MDA-MB-231 cells were transfected with a non-targeting siRNA (siCTRL), siRNA targeting syntenin (siSYNT_03 and siSYNT_04), or siRNA targeting SDC1 and SDC4 (siSDC1+4). (**A**) Representative western blot of biotinylated cell surface proteins, MESNA-treated fractions (control for surface biotin removal), and fractions corresponding to 30 minutes of endocytosis. MT1-MMP specific antibody was used. GAPDH in total cell lysates is used as a loading control. (**B**) Dot plot represents the mean signal of MT1-MMP cell surface levels normalized by GAPDH from cell lysate, from five independent experiments for siSYNT_03, six for siSYNT_04, and four for siSDC1+4. (**C**) Dot plot represents the fraction of MT1-MMP internalized from the cell surface, normalized to its surface levels, from four independent experiments. Dot plots show the number (**D**) and the size (**E**) of sEV purified from the culture medium, analyzed using Nanosight technology (NTA). (**F**) Representative western blot of cell lysates and of the same number of sEV. Specific antibodies were used to detect ALIX, MT1-MMP, syntenin, flotillin, CD9, and SDC1-CTF. Red Ponceau staining was used as a loading control. (**G-H**) Dot plots represent the mean signal of MT1-MMP protein levels in cell lysates (**G**) and in sEV fractions (**H**), normalized to Red Ponceau (Red P) staining. The Paired Student’s t-test with Welsh correction was used for statistical analysis. *** p ≤ 0.001, ** p ≤ 0.01, * p ≤ 0.05, ns: not significant.

Given the importance of syntenin and SDC in the biogenesis of sEV from endosomal origin (Baietti et al., 2012), we hypothesize that they may modulate MT1-MMP sorting in sEV and its subsequent extracellular functions. Here, we refer to sEV rather than sEV from endosomal origin, in accordance with the MISEV2023 guidelines (Welsh et al., 2024), as our purification method does not allow us to definitively determine the origin of the isolated vesicles. The analysis of sEV isolated from conditioned media by ultracentrifugation (concentrated in sEV from endosomal origin) showed a 20% reduction in sEV number upon depletion of syntenin or SDC in MDA-MB-231 TNBC cells (Figure 4D), while sEV size remained unchanged (Figure 4E). Furthermore, key regulators of sEV biogenesis, such as ALIX (Figure S1A) and SDC (Figure S1E) (predominantly found as SDC-CTF, corresponding to the membrane-associated remnant of protease-cleaved SDC), whose loading in sEV is controlled by syntenin and SDC, were decreased in sEV fraction (Figure 4F) (Baietti et al., 2012), confirming that the sEV biogenesis pathway controlled by syntenin and SDC is functional in MDA-MB-231 cells. The analysis of sEV-associated MT1-MMP revealed a striking 50% decrease in MT1-MMP incorporation upon syntenin or syndecan depletion, with a slight increase of total MT1-MMP protein levels (Figure 4G-H). Importantly, the levels of canonical syntenin-independent sEV markers (flotillin (Figure S1B), CD9 (Figure S1D)) remained unchanged across conditions, further supporting the notion that syntenin and SDC selectively regulate MT1-MMP loading into sEV.

These results establish syntenin and SDC as key regulators of MT1-MMP sorting into sEV, without influencing its delivery to the cell surface and its endocytosis.

### sEV contribute to ECM degradation *via* MT1-MMP activity in a syntenin-dependent manner

Since MT1-MMP-containing sEV were efficiently secreted from MDA-MB-231 cells, we next investigated whether these vesicles had the capacity to degrade the ECM. We isolated sEV from culture medium by immunocapture using CD9, CD63, and CD81 antibodies-coated beads and seeded them associated to the beads onto a fluorescently labelled gelatin for 24 hours. sEV derived from non-targeting siRNA (siCTRL) transfected cells exhibit proteolytic activity, forming distinct degradation zones that colocalized with the exosomal marker CD9 (Figure 5B). A previous study demonstrated that the use of a general inhibitor of MMP (GM6001) inhibited the capacity of sEV to degrade the matrix (Hakulinen et al., 2008). Here, we show that depletion of MT1-MMP using a specific siRNA (siMT1-MMP) in sEV-producing MDA-MB-231 cells strongly abolished sEV-associated degradation zones (Figure 5A), demonstrating that MT1-MMP is essential for ECM degradation mediated by sEV.

**Figure 5:**
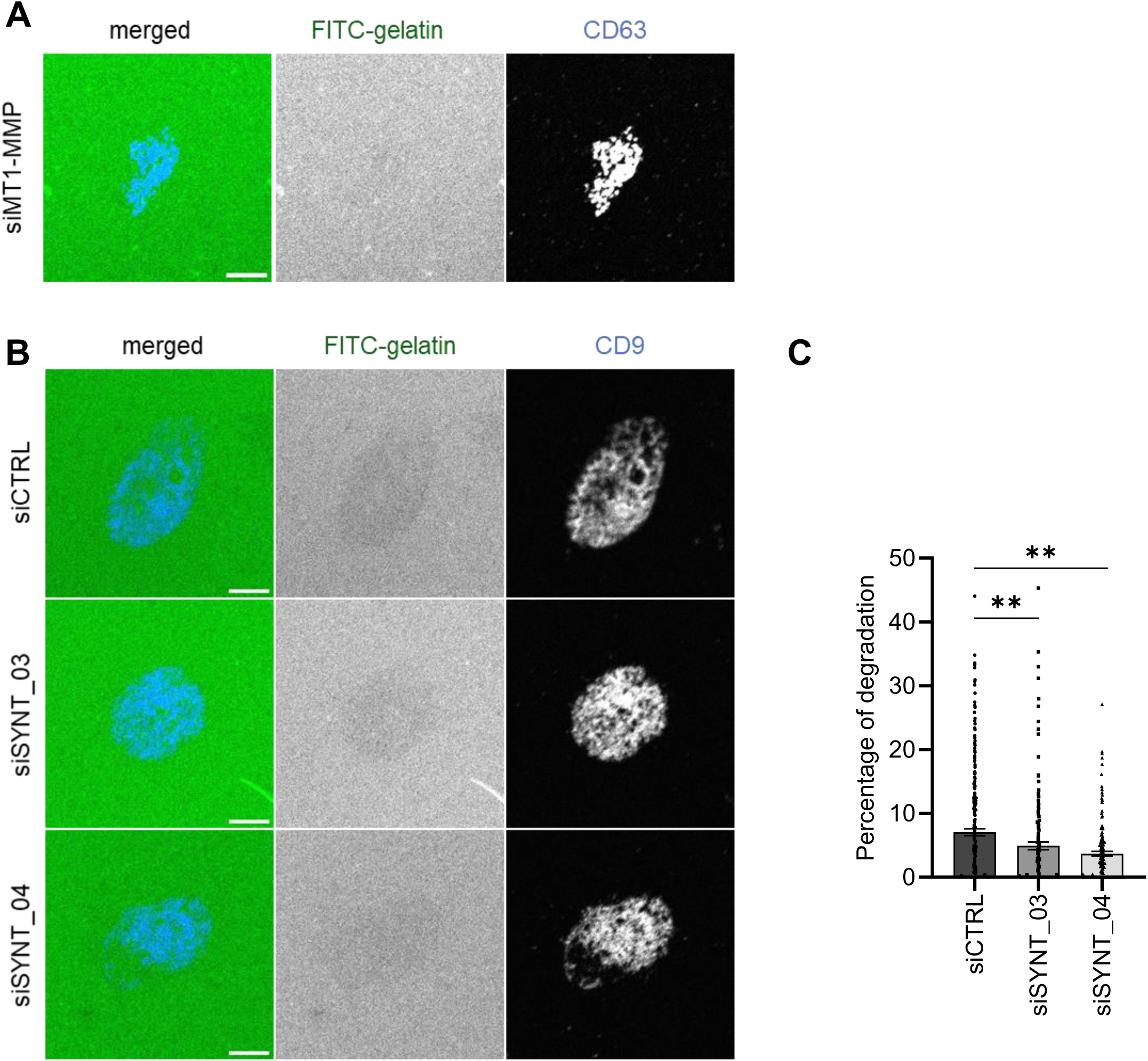
sEV contribute to ECM degradation through MT1-MMP activity, a process dependent on syntenin. Representative images of sEV produced by MDA-MB-231 cells transfected with a (**A**) siRNA targeting MT1-MMP, (**B**) non-targeting siRNA (siCTRL) or siRNA targeting syntenin (siSYNT_03 and siSYNT_04), isolated by immunocapture, and seeded on fluorescently-labeled gelatin (FITC-gelatin) for 24 hours. sEV were fixed and stained with an anti-CD63 or anti-CD9 antibody. (**C**) Dot plot represents the percentage of gelatin degradation determined from at least 60 sEV-enriched zones ± SEM from three independent experiments. The Mann-Whitney test was used for statistical analysis. **** p ≤ 0.0001, *** p ≤ 0.001, ** p ≤ 0.01, * p ≤ 0.05, ns: not significant.

Given that syntenin is responsible for MT1-MMP loading into sEV, we next examined the impact of syntenin on the degradative potential of sEV. sEV derived from syntenin-depleted cells exhibited a significantly reduced percentage of ECM degradation (Figure 5B-C). These findings indicate that MT1-MMP incorporation into sEV is functionally relevant for ECM proteolysis controlled by syntenin.

## Discussion

Invasive cancer progression relies on the ability of tumour cells to remodel the ECM, a process that is tightly regulated by proteolytic degradation of ECM components by MMP (Azadi and Tafazzoli Shadpour, 2020; Winkler et al., 2020). In this study, we clarified how syntenin, SDC, and MT1-MMP interact to control ECM degradation, in particular through invadopodia-dependent and sEV-mediated mechanisms.

In line with the described pro-invasive function of syntenin, this study describes syntenin has a positive regulator of ECM degradation. Indeed, syntenin controls the formation and the activity of invadopodia developed by cancer cells. The pronounced impact of syntenin depletion on the number of mature invadopodia suggests that syntenin may be directly implicated in the delivery of MT1-MMP to precursor invadopodia, thereby enabling their enzymatic activation. However, we cannot rule out that syntenin effect is indirect through regulation of the EMT process that is linked to invadopodia formation (Eckert et al., 2011) as syntenin has been described to regulate EMT, notably through control of the TGF-β/Smad signalling pathway (Hwangbo et al., 2016). It can also be a consequence of impaired sEV secretion as we will discuss it later.

Although syntenin and SDC are known to cooperate in the regulation of sEV from endosomal origin biogenesis, our findings reveal that they exhibit opposing roles in ECM degradation by cells. Indeed, taken together, SDC1 and SDC4 seem to limit ECM degradation by cells (Figure 1). Furthermore, while syntenin seems to have an impact on invadopodia formation and maturation, our quantitative analysis suggests that SDC would affect the dynamics of invadopodia assembly and disassembly, rather than their formation and maturation. This divergence highlights that proteins classically viewed as partners in one cellular context may engage distinct roles in another cellular context. Mechanistically, this functional divergence suggests the existence of a context-dependent regulatory mechanism allowing these proteins to perform distinct roles depending on the cellular signals present. It is also worth noting that, in our MDA-MB-231 cellular model and in other breast cancer cell lines, a dichotomy has been reported between SDC1 and SDC4. Indeed, SDC1 has been described to act as an anti-migratory/invasive factor (Ibrahim et al., 2012), whereas SDC4 has been associated with pro-migratory/invasive functions (Onyeisi et al., 2021). Our results suggest that, in the context of ECM degradation, the anti-migratory/invasive function of SDC1 may override the promigratory/invasive function of SDC4. The individual contribution of SDC to ECM proteolysis warrants further investigation. Importantly, for other key processes of tumour progression, such as induction of angiogenesis or acquisition of a cancer stemness phenotype, SDC1 displays pro-tumorigenic functions (Ibrahim et al., 2013; Nassar et al., 2021). However, the role of SDC in cancer progression is highly tissue-specific, depending not only on their expression levels in cancer and stromal cells but also their localization (e.g. shed extracellular vs. membrane-bound forms) (Szatmári et al., 2015).

Classically, interactions between PDZ domain-containing proteins and their partners occur through the recognition of a PDZ BM located at the C-terminus of the partner protein (Lee and Zheng, 2010). In some cases, the PDZ BM has been described to be internal (Hillier et al., 1999; Penkert et al., 2004). Our results reveal that, although the ICD of MT1-MMP bears a C-terminal located PDZ BM, this motif is not involved in its interaction with syntenin, a PDZ domain-containing protein. Indeed, deletion of the PDZ BM (DKV) in the ICD of MT1-MMP does not disrupt the interaction. Our data indicate that this interaction is mediated instead by a PRR motif within the ICD of MT1-MMP (Figure 3). The PRR motif does not fulfil the typical characteristics of a PDZ BM, which classically features a hydrophobic amino acid at the C-terminal position (p0), and, depending on the PDZ BM class, either a hydrophobic, T/S, or D/E at the p-2 position, pointing to a non-canonical but specific mode of binding between the region of syntenin containing its two PDZ domains and an alternative functional region of MT1-MMP. As previously reported (Lee and Zheng, 2010; Ye and Zhang, 2013), this illustrates the structural and functional flexibility of the PDZ domain, allowing it to engage with alternative motifs depending on the available partners. The interface involved in syntenin needs to be investigated.

In addition, syntenin interacts with SDC through PDZ-dependent interactions (Grootjans et al., 2000). This interaction is essential for the biogenesis of sEV from endosomal origin and the loading of specific cargoes (Baietti et al., 2012). MT1-MMP and SDC have been previously functionally linked. Indeed, MT1-MMP induces the extracellular cleavage of SDC and this impacts cell adhesion and migration (Endo et al., 2003; Manon-Jensen et al., 2013). Here, we describe that both syntenin and SDC are required for the efficient loading of MT1-MMP into sEV (Figure 4), suggesting that they may act in a complementary or concerted manner as part of a shared sorting mechanism. We therefore hypothesize that the interaction between syntenin and MT1-MMP may be at least partially indirect, or facilitated by SDC, within a tripartite complex. This complex could form transiently or partially and involve both intracellular interactions (between syntenin, SDC and MT1-MMP) and extracellular interactions (between MT1-MMP and SDC), orchestrating the selective sorting of MT1-MMP into sEV. Though not yet demonstrated, such an arrangement would be compatible with a multi-layered regulatory mechanism involving structural recognition, post-translational modifications (cleavage), and specialized functions in sEV biogenesis. This model highlights the complexity of PDZ domain-mediated interaction networks, which may go beyond traditional binary schemes to incorporate more flexible mechanisms, including direct and indirect interactions adaptable to subcellular context.

Our findings show that syntenin interacts specifically with MT1-MMP through the PRR motif in its ICD (Figure 3). This motif has been previously identified as a binding site for MTCBP1 a protein known for its anti-tumoral effects (Qiang et al., 2018; Uekita et al., 2004). The convergence of two functionally opposing proteins at the same interaction site suggests a competitive mechanism for binding to MT1-MMP. Such competition could fine-tune cellular signalling by shifting the balance between pro- and anti-tumour responses, depending on which partner is recruited to the PRR motif. In the tumor context, an increased expression of syntenin, or a higher binding affinity for the PRR motif, could favour its interaction at the expense of MTCBP1, potentially disrupting the intracellular signalling balance. This imbalance may contribute to tumour progression by impairing protective pathways. From a therapeutic perspective, this interaction interface presents an interesting target. Identifying molecules that specifically modulate the syntenin-MT1-MMP interaction, without disrupting MTCBP1 binding, could help restore protective signalling in tumour cells. Understanding the structural and functional determinants governing the recognition of the PRR motif could open the way for targeted modulation strategies. Overall, our findings highlight the PRR motif as a crucial platform for integrating antagonistic cellular signals, and its regulation could be key to understanding tumour progression and developing therapeutic strategies aimed at disrupting this balance.

Late endosomes serve as critical turntable for the sorting, recycling, or degradation of internalized proteins, orchestrating key trafficking decisions within the cell (Scott et al., 2014). Within these compartments, we observed a preferential colocalization of syntenin and MT1-MMP (Figure 2). This spatial proximity supports a functional interaction between syntenin and MT1-MMP in late endosomes, that we demonstrate to contribute to the sorting of MT1-MMP into sEV from endosomal origin (Figure 4). By promoting the loading of MT1-MMP into sEV, syntenin could not only protect MT1-MMP from lysosomal degradation but also enable its sustained proteolytic activity in the tumour microenvironment. Normally, lysosomes target internalized proteins for degradation after their entry into late endosomes (Pryor and Luzio, 2009), however, the incorporation of MT1-MMP into sEV could bypass this pathway, providing a protective compartment for the enzyme, allowing it to remain functional and extend its half-life.

The role of MT1-MMP in ECM degradation is central to cancer cell invasion, as this metalloproteinase is known to cleave key structural components of the ECM (Itoh, 2015), facilitating tissue penetration. In the context of sEV, our results demonstrate that MT1-MMP becomes actively involved in ECM remodelling by acting as a carrier of degradative potential (Figure 5). This proves that sEV loaded with MT1-MMP directly contribute to ECM degradation, enabling cancer cells to modify the extracellular environment even at a distance from the primary tumour site. In this way, the loading of MT1-MMP into sEV represents a dynamic mechanism by which tumour cells can enhance their invasive capacity through the coordinated proteolysis of the ECM, without requiring direct cell contact.

Considering the established role of syntenin and SDC in sEV biogenesis (Baietti et al., 2012), along with our findings demonstrating their involvement in MT1-MMP loading into sEV (Figure 4), these components appear essential for maintaining the balance between degradation and protective transport of this critical protease in the tumour microenvironment. As MT1-MMP-loaded sEV contribute to ECM remodelling and cancer cell invasion, syntenin likely modulates not only the stability of MT1-MMP but also its ability to participate in matrix degradation (Figure 5). This regulation of MT1-MMP in the context of sEV biogenesis could be pivotal in controlling tumour progression, where the availability of active proteases in the extracellular space is essential for driving invasive behaviour. By hijacking this pathway, tumour cells may be able to enhance ECM degradation at multiple sites, facilitating metastasis and altering the tumour microenvironment to favour invasion and dissemination.

It is now established that invadopodia and sEV are functionally connected. Firstly, invadopodia have been described to represent a preferential site of MVB fusion (Hoshino et al., 2013). Moreover, inhibition of sEV secretion impairs both the formation and activity of invadopodia, and *vice versa* (Hoshino et al., 2013). In this context, our results showing that syntenin controls both invadopodia formation and maturation (Figure 1) and sEV secretion (Figure 4), suggest that syntenin may act directly on invadopodia formation, but also indirectly through regulation of sEV secretion. This hypothesis is consistent with the notion that sEV contribute to the local supply of proteolytic enzymes, such as MT1-MMP, and modulate the tumour microenvironment. Therefore, the observed negative impact of syntenin depletion on ECM degradation could result from impaired sEV-mediated signalling or cargo delivery, such as MT1-MMP which loading is strongly affected by syntenin (Figure 4). Further studies will be required to determine whether the effects of syntenin are exclusively mediated *via* sEV, or whether a direct sEV-independent regulation of invadopodia also occurs. Interestingly, it has been shown that exosomes carrying MT1-MMP can be retained at the cell surface *via* BST2/tetherin, enhancing pericellular matrix degradation and tumour invasion (Palmulli et al., 2025). It would be interesting to assess whether syntenin depletion also affects sEV retention at the plasma membrane, potentially influencing their local activity in invasion processes.

The interaction between syntenin and MT1-MMP represents a key regulatory node in the invasive behaviour of tumour cells, particularly through the biogenesis and molecular composition of sEV. Targeting this interaction may provide a promising strategy to impair tumour invasion by limiting the trafficking of MT1-MMP into sEV, thereby reducing their ability to remodel the ECM. Conversely, an opposite strategy can be envisioned in a therapeutic context: harnessing the invasive potential of MT1-MMP by enhancing its loading into engineered sEV designed for drug delivery. By increasing MT1-MMP enrichment in these vesicles, it may be possible to improve their ability to penetrate dense tumour stroma, which often acts as a major physical barrier to treatment efficacy. This dual approach, either inhibiting the syntenin/MT1-MMP interaction to block tumour invasion or exploiting it to potentiate stromal infiltration by therapeutic sEV, highlights the versatility of this pathway and its potential for context-dependent therapeutic applications.

## Materials and methods

### Cell lines

MDA-MB-231 TNBC-derived cells were maintained in Dulbecco’s Modified Eagle Medium (Thermo Fisher, USA) supplemented with 10% fetal bovine serum (FBS) (Sigma, Life Science, USA) at 37 °C in a humidified atmosphere containing 5% CO₂. Mycoplasma contamination was regularly monitored using the Mycoplasmacheck qPCR-based detection service (Eurofins genomics, Germany). The cultured cells were split every 3-4 days, maintaining an exponential growth.

### Sequences of siRNA and transfection

RNAi oligonucleotides targeting human syntenin-1 (or SDCBP-1) (siSYNT_03 and siSYNT_04), syndecan-1 (siSDC1_04), syndecan-4 (siSDC4_01), and a SMARTpool targeting human MT1-MMP (siMT1-MMP), as well as a nontargeting control RNAi (siCTRL), were obtained from Dharmacon Inc (USA).

For RNAi experiments, suspended cells were transfected with 20 nM RNAi using Lipofectamine RNAiMAX reagent (Life Technologies, USA). Five hours post-transfection, the medium was replaced with a transfection-free medium. Cells were analysed 48 or 72 hours after RNAi treatment.

### Antibodies

Antibodies directed against syntenin and ALIX were described earlier (Baietti et al., 2012). CD63 and CD9 antibodies were provided by E. Rubinstein (Charrin et al., 2001) (Université Paris-Sud UMRS_935, Villejuif). Other antibodies were from commercial sources and were used as recommended by the manufacturers. Antibodies directed against MT1-MMP rabbit (ab51074) and syntenin (ab133267) were from Abcam (UK), MT1-MMP mouse (MAB3328) and cortactin (05-180-I-25UL) were from Millipore (USA), syndecan-1 intracellular domain (D4H7H) was from Cell Signaling (USA), syndecan-4 intracellular domain (PAB9045) was from Abnova (Taiwan), EEA1 (sc-6415) and LAMP2 (sc-18822) were from Santa Cruz (USA), anti-GM130 (610822) and anti-flotillin-1 (610820) were from BD Biosciences (USA), TKS-5 (NBP1-90454) was from Novus biologicals (USA) and GAPDH (10494-1-AP) was from Proteintech (USA). The HRP linked secondary antibodies directed against mouse (NA931) and rabbit (NA934) were from Cytiva (USA) and the Alexa Fluor 405-, Alexa Fluor 488-, Alexa Fluor 555-, Alexa Fluor 647-conjugated secondary antibodies were from Invitrogen (USA).

### sEV isolation and total cell lysates

sEV fractions were collected from equivalent amounts of culture medium conditioned by equivalent numbers of seeded cells (4×10^6^ cells for dUC and 1.5×10^7^ cells for immunocapture). Following the required treatments, cells were refreshed with media containing 10% sEV-depleted FBS, which was depleted of sEV by overnight centrifugation at 110,000 x g. Conditioned media were collected after 16 hours. Lysates from corresponding cultures were prepared by scraping cells in lysis buffer (30 mM Tris–HCl pH 7.4, 150 mM NaCl, 1% NP-40, protease inhibitors), then lysed for 30 minutes and cleared by centrifugation at 10,000 x g for 10 minutes at 4°C. Protein concentration was quantified using a BCA assay (Thermo Scientific Pierce, USA).

#### sEV isolation by differential ultra-centrifugation (dUC)

For comparative analyses in loss-of-function studies, sEV were isolated through differential centrifugation at 4°C as follows: 10 minutes at 1,500 x g to remove cells and large debris, 30 minutes at 9,000 x rpm to eliminate large extracellular vesicles (corresponding mainly to microvesicles), and 1.5 hours at 29,000 x rpm to pellet sEV (corresponding mainly to exosomes). A final wash with PBS was performed, followed by centrifugation at 55,000 x rpm for 1 hour to remove soluble serum components and secreted proteins. Cell lysates normalized to the amount of protein after a BCA assay and sEV lysates from equivalent number of sEV were subsequently analyzed by Western Blot.

#### sEV isolation by immunocapture

sEV isolation was performed by positive selection using MicroBeads recognizing the tetraspanin proteins CD9, CD63, and CD81 (EV Isolation Kit Pan, human, Miltenyi Biotec, Bergisch-Gladbach, Germany). The culture medium was centrifuged for 10 minutes at 1,500 x g to remove cells and large debris, followed by 30 minutes at 9,000 x rpm to eliminate larger extracellular vesicles (microvesicles). The cleared supernatant was concentrated by ultrafiltration at 5,000 x rpm using Amicon Ultra-15 Centrifugal Filter Units – 10 kDa (Millipore, Billerica, MA, USA) until 2 mL was obtained. The concentrated supernatant was incubated with the beads. The sEV associated to beads were then selected and eluted according to the manufacturer’s protocol.

### Nanoparticle Tracking Analysis (NTA)

sEV collected by ultracentrifugation were resuspended in PBS and analyzed at a 1/50 dilution using a Nanosight NS-300 instrument (Malvern Panalytical, UK). For each sample, three 60-second videos were recorded at 25°C, with 20–60 particles per frame, to determine mean particle size and concentration.

### Western Blot

Proteins were heat-denatured in loading buffer (Tris-HCl 250 mM pH 6.8, SDS 10%, Glycerol 25%, Bromophenol Blue), fractionated by SDS-PAGE using NuPage 4-12% Bis-Tris Gel (Invitrogen, USA), and electrotransferred onto nitrocellulose membranes. Membranes were stained with Ponceau Red and immunoblotted with the indicated primary antibodies, followed by HRP-conjugated secondary antibodies. Signals were detected using the Amersham ECL Prime and Select Western Blotting Detection Reagent (GE Healthcare, France) and quantified by densitometric analysis with ImageJ software.

### Immunofluorescence staining and confocal microscopy

Glass coverslips were pre-coated with 5 µg/cm² collagen I (Life Technologies, USA) at 37°C for at least 30 minutes. MDA-MB-231 cells were then seeded onto the coverslips and incubated for 24 hours. After incubation, cells were fixed with 4% paraformaldehyde for 20 minutes at room temperature (RT), permeabilized with 0.1% Triton X-100 for 10 minutes, and blocked with 1% BSA for 1 hour. Cells were then stained with the respective primary and secondary antibodies, mounted in ProLong™ Diamond Antifade Mountant (Invitrogen, USA) containing DAPI (Merck, Germany), and imaged using a confocal NIKON AX (NIS-Elements) equipped with the appropriate lasers and a 60x objective. Confocal images were analysed using Image J software. Colocalization between MT1-MMP and syntenin was assessed using the JaCoP (Just Another Colocalization Plugin) plugin, which calculates the Pearson’s correlation coefficient between the two fluorescence channels. Analyses were performed on manually selected regions of interest corresponding to individual cells. For compartment specific quantification, a custom ImageJ macro was used to generate binary masks from fluorescence channels. The masks were used to segment intracellular compartments, and the proportion of each compartment area occupied by the colocalized MT1-MMP and syntenin signals was subsequently measured.

### Gelatin matrix degradation assay and invadopodia/degradation quantification

FITC gelatin-coated coverslips were prepared by incubating coverslips with 50 μg/ml poly-D-lysine (Merck, Germany) for 20 minutes at 37 °C, followed by 0.5% glutaraldehyde (Merck, Germany) for 15 minutes at RT. The coverslips were then inverted onto a drop of 0.2% gelatin mixed with fluorescently labeled gelatin (Oregon Green 488, 10:1) for 10 minutes at RT. The gelatin matrix was quenched with 5 mg/ml sodium borohydride (Merck, Germany) for 5 minutes at RT and dehydrated in 70% ethanol for storage.

For cell degradation assays, 48 hours post-transfection, MDA-MB-231 cells were detached using 1 mM PBS-EDTA, and 50000 cells were plated on gelatin-coated coverslips, centrifuged for 2 minutes at 900 x rpm to synchronize cell adhesion and incubated for 5 hours at 37°C.

Immunofluorescence protocol was performed. Primary antibodies against TKS5 and cortactin, followed by Alexa Fluor 555- and Alexa Fluor 405-conjugated secondary antibodies, respectively, were used for immunolabeling. Images were acquired using a Zeiss ApoTome structured light microscope equipped with a 63 x 1.4 Plan-Apochromat objective. The percentage of cells undergoing degradation was assessed by counting 10 random fields at 63x magnification. The total number of cells and the number of cells displaying matrix degradation were recorded to calculate the percentage of degrading cells relative to the total cell population. Twenty-five cells per condition per experiment were analysed to quantify invadopodia parameters using a custom Fiji macro (Chanez et al., 2021; Thuault et al., 2020). A Gaussian filter (sigma = 80) was applied to reduce background noise. Cell area was measured in the red channel (TKS5) by manually delineating the cell contour. Cortactin (blue channel), TKS5 (red channel), and degradation areas (green channel) were detected following manual thresholding, binarization and morphological correction (hole filling). Positive regions were quantified using the *Analyse Particles* tool with measurements including number, area, mean pixel intensity. Channels were merged to generate composite and grayscale images for global signal distribution analysis. Colocalizations were assessed using binary channel combination with the *Image Calculator* (AND), followed by particles analyses. The number of total invadopodia is defined by the co-localization of TKS5 and cortactin, and the number of mature invadopodia by the co-localization of TKS5, cortactin, and degradation foci.

For sEV degradation assays, half of the sEV previously isolated using immunocapture were deposited on gelatin-coated coverslips, centrifuged for 2 minutes at 900 x rpm and incubated for 24 hours at 37°C. sEV were then fixed with 4% paraformaldehyde, permeabilized with 0.1% Triton X-100, and blocked with 1% BSA. Immunolabeling was performed using primary antibodies against CD9, followed by Alexa Fluor 405-conjugated secondary antibodies. Images were acquired using a confocal NIKON AX (NIS-Elements). For each condition and experiment, ten images were analysed. Image analysis was performed using a custom macro in Image J to quantify ECM degradation associated to sEV signal. A Gaussian blur filter (sigma = 2) was applied to the sEV channel (CD9 staining) to reduce background noise. A fixed threshold value was applied to sEV channel to identify regions enriched in sEV. These regions were binarized, and particles larger than 3 pixels^2^ were detected using the “Analyse particles” function. This step generated a set of ROIs corresponding to sEV-enriched zones. The mean fluorescence intensity of fluorescent gelatin was then measured globally and within each ROI to quantify the degree of gelatin degradation. These data were then processed to calculate the percentage of gelatin degradation per individual sEV-enriched zone, based on the mean field intensity and the intensity within the degradation area.

### Cell Surface Protein Biotinylation and Endocytosis

Cell surface proteins were biotinylated twice with 0.2 mg/mL reducible sulfo-NHS-SS-biotin (Thermo Scientific Pierce, USA) at 4°C for 2 x 30 minutes. Following biotinylation, cells were washed three times with PBS Ca/MgCl_2_ (0.5 mM MgCl_2_ + 1 mM CaCl_2_), and unreacted biotin was quenched 5 minutes using a 50 mM Tris-HCl (pH 8.0) solution.

For endocytosis experiments, after biotinylation, cells were incubated at 37°C for 30 minutes in culture medium to allow internalization of cell surface proteins. The process was stopped by a cold PBS Ca/MgCl_2_ wash, and remaining cell surface biotin was removed by 2 x 20 minutes treatment with 50 mM MesNa (sodium 2-mercaptoethanesulfonate, Sigma, USA) solubilized in 50 mM Tris pH 8.6, 50 mM NaCl, 1 mM EDTA, 0.2% BSA, and 1 mM CaCl_2_. For negative control, before protein endocytosis, cell surface biotin was removed by 2 x 20 minutes treatment with 50 mM MesNa.

Cells were then lysed for 30 minutes in lysis buffer (30 mM Tris–HCl pH 7.4, 150 mM NaCl, 1% NP-40, protease inhibitors). The lysates were cleared by centrifugation at 10,000 x g for 10 minutes at 4°C, and the resulting supernatants were incubated with streptavidin beads (Merck, USA) for 1.5 hours at 4°C under continuous mixing. After incubation, the beads were washed three times with PBS, resuspended in 2X loading buffer, and boiled at 95°C for 10 minutes. Biotinylated proteins were then detected by western blot as described above.

### Plasmid constructs and biotinylated peptides

GST-Syntenin Tandem + Cter WT, GST-Syntenin Tandem PDZ1^K119A^ ^+ SDK^ ^to^ ^HEQ^ + Cter, GST-Syntenin Tandem PDZ2^K203A^ ^+ KDS^ ^to^ ^SHE^ + Cter and GST-Syntenin Tandem PDZ1^K119A^ ^+ SDK^ ^to^ ^HEQ^ +PDZ2^K203A^ ^+ KDS^ ^to^ ^SHE^ + Cter cDNAs were described in Grootjans et al, 2000 (Grootjans et al., 2000). All constructs were verified by DNA sequencing (Eurofins Genomics, Germany). The biotinylated peptides were designed and ordered from Covalab (France).

### Recombinant protein production and purification

#### Plasmid transformation in BL21 Bacteria

Plasmids were transformed in chemically competent BL21 bacteria (Thermofisher Scientific, USA), following manufacturer recommendations.

#### Colony Culture and Induction of Protein Expression

A single transformed colony was selected and inoculated into 15 mL of Terrific Broth medium (Tryptone 1.2%, yeast extract 2.4%, glycerol 0.5%) containing the appropriate antibiotic (50 µg/mL), followed by overnight incubation at 37°C with shaking. The culture was then diluted into 500 mL of TB medium and grown at 37°C until the optical density at 600 nm reached 0.4 - 0.6. Protein expression was then induced by adding 1 mM IPTG (Generon, UK), and incubation was continued at 30°C with shaking for 5 hours. After centrifugation (8000 x rpm, 4°C, 20 minutes), the bacterial pellet was stored at -20°C for subsequent protein purification.

#### Recombinant Protein Purification

Bacterial pellets were resuspended in lysis buffer (25mM Tris pH 8, 150mM NaCl, 1X protease inhibitor cocktail (Life technologies, Pierce, USA), DNase I (1µg/mL), and lysozyme (0.4 mg/mL) (Sigma, USA)) and incubated on ice for 30 minutes with intermittent vortexing. Cell lysis was performed by sonication using a Vibra-Cell 750 (Sonics & Materials, UK) for 10 cycles of 10 seconds at 35%, followed by 4 cycles of 40 seconds in pulse mode (2s ON/1s OFF). After centrifugation at 12000 x g for 45 minutes at 4°C, the supernatant was filtered (0.45 µm) and subjected to purification using a GSTrap FF Column (Cytiva, USA) on an ÄKTA Prime system (GE Healthcare, France). After washing the column with equilibration buffer (25 mM Tris pH 8, 150 mM NaCl), the fusion protein was eluted from the column with a gradient of reduced glutathione (0-50 mM) (Sigma, USA). The purity of the fusion proteins was analysed by SDS-PAGE followed by Coomassie Blue staining.

### Surface Plasmon Resonance (SPR) experiments

All SPR experiments were performed at 25°C using a BIAcore T200 instrument (GE Healthcare, France). Biotinylated peptides were immobilized on a streptavidin-sensor chip (Cytiva, USA) at a density of 1000 resonance units (RU). A blank-immobilized reference channel was used for background subtraction. GST fusion protein (analytes) were perfused at a 30 μL/min flow rate in running buffer (25 mM Tris-HCl, 150 mM NaCl, 0.005 % Tween 20). Analytes were injected at varying concentrations (in μM) for 300 s or 120 s, allowing sensorgrams to reach equilibrium, followed by a 120 s dissociation phase. Binding was monitored as the difference between the response from the functionalized surface and the reference surface. Between runs, sensor chip surfaces were regenerated with short pulses of 50 mM NaOH and 1 M NaCl at a 30 μL/min flow rate to ensure complete removal of bound analytes.

### Statistical analysis

All statistical analyses were performed using GraphPad Prism software (USA). Depending on the dataset, statistical significance was determined using the Mann-Whitney test, or paired Student’s t-test with Welsh correction. Graphs were plotted using GraphPad Prism software, showing the mean ± SEM. The significance thresholds were as follow: p > 0.05 (ns, non-significant), *p ≤ 0.05, **p ≤ 0.01, ***p ≤ 0.001, and ****p ≤ 0.0001. The number of samples, images, and experiments used for quantification are provided in the respective figure legends, with n representing the number of cells and N indicating the number of experiments.

## Supporting information

Huber et al Supplementary Information

## Acknowledgments

We thank M. Rodriguez (CRCM Microscopy Platform) for her assistance with image acquisition and the development of analyses macros. We are grateful to S. Granjeaud (CRCM Bioinformatic Platform) for valuable help with statistical analyses.

