## Supplementary material for "Syntenin orchestrates matrix degradation by controlling MT1-MMP secretion in small extracellular vesicles and invadopodia formation": Huber et al Supplementary Information

**<u>Figure S1:</u> Protein enrichment in sEV following syntenin and syndecan depletion**

MDA-MB-231 cells were transfected with a non-targeting siRNA (siCTRL), siRNA targeting syntenin (siSYNT_03 and siSYNT_04), or siRNA targeting SDC1 and SDC4 (siSDC1+4). Dot plots represent the mean signal of ALIX (**A**), Flotillin (**B**), Syntenin (**C**), CD9 (**D**) and SDC1-CTF (**E**) protein levels in cell lysates and in sEV fractions, normalized to Red Ponceau (Red P) staining. The Paired Student's t-test with Welsh correction was used for statistical analysis. **** $p \leq 0.0001$, *** $p \leq 0.001$, ** $p \leq 0.01$, * $p \leq 0.05$, ns: not significant.

**Figure S1**

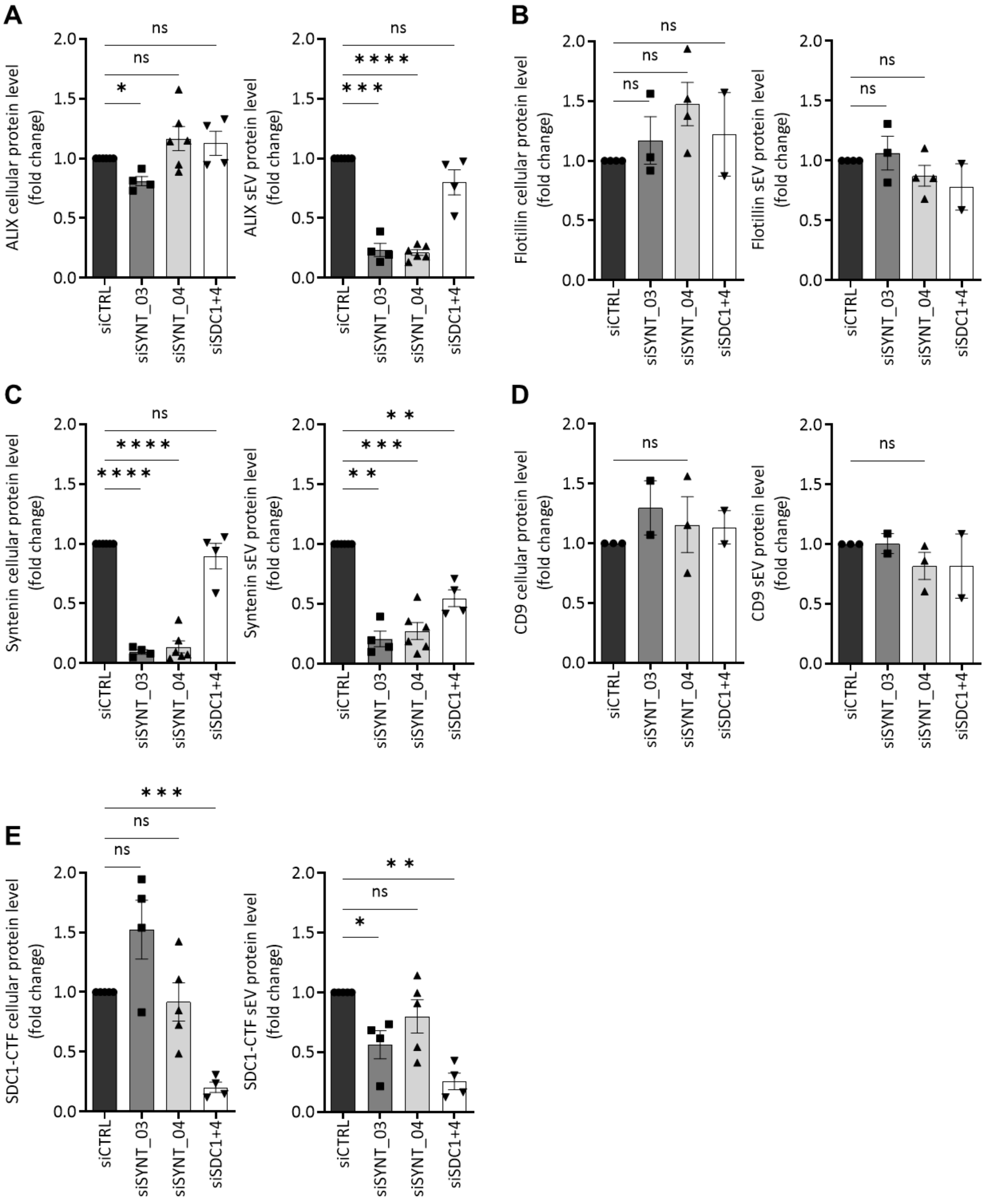
A
ALIX cellular protein level
(fold change)
ALIX sEV protein level
(fold change)
B
Flotillin cellular protein level
(fold change)
Flotillin sEV protein level
(fold change)
C
Syntenin cellular protein level
(fold change)
Syntenin sEV protein level
(fold change)
D
CD9 cellular protein level
(fold change)
CD9 sEV protein level
(fold change)
E
SDC1-CTF cellular protein level
(fold change)
SDC1-CTF sEV protein level
(fold change)
0.0
0.5
1.0
1.5
2.0
siCTRL
siSYNT_03
siSYNT_04
siSDC1+4
ns
*
**
***
****
